# Two parallel pathways are required for ultrasound-evoked behavioral changes in *Caenorhabditis elegans*

**DOI:** 10.1101/2021.10.29.466533

**Authors:** Uri Magaram, Connor Weiss, Aditya Vasan, Kirthi C. Reddy, James Friend, Sreekanth H. Chalasani

**Affiliations:** Neurosciences Graduate Program, University of California San Diego, La Jolla, CA 92093; Molecular Neurobiology Laboratory, The Salk Institute for Biological Studies, La Jolla, CA 92037; Medically Advanced Devices Laboratory, Department of Mechanical and Aerospace Engineering, Jacobs School of Engineering and Department of Surgery, School of Medicine, University of California San Diego, La Jolla, CA 92093

**Keywords:** Ultrasound, *C. elegans*, reversals, *trp-4*, *mec-4*, Sonogenetics, parallel pathways

## Abstract

Ultrasound has been shown to affect the function of both neurons and non-neuronal cells. However, the underlying molecular machinery has been poorly understood. Here, we show that at least two mechanosensitive proteins act in parallel to generate *C. elegans* behavioral responses to ultrasound stimuli. We first show that these animals generate reversals in response to a single 10 msec pulse from a 2.25 MHz ultrasound transducer. Next, we show that the pore-forming subunit of the mechanosensitive channel TRP-4, and a DEG/ENaC/ASIC ion channel MEC-4, are both required for this ultrasound-evoked reversal response. Further, the *trp-4 mec-4* double mutant shows a stronger behavioral deficit compared to either single mutant. Finally, overexpressing TRP-4 in specific chemosensory neurons can rescue the ultrasound-triggered behavioral deficit in the *mec-4* null mutant, suggesting that these two pathways act in parallel. Together, we demonstrate that multiple mechanosensitive proteins likely cooperate to transform ultrasound stimuli into behavioral changes.

## Introduction

Ultrasound has been shown to modify neuronal activity in number of animal models including humans [1, 2]. However, the direction of this action is somewhat controversial with some reporting activation [3-7], while others demonstrate inhibition [2, 8-10]. Moreover, the underlying mechanisms for ultrasound action on neuronal membranes have been suggested to include thermal [11-13], mechanical (direct or via cavitation [14-16]) or a combination of two [17]. Additionally, ultrasound neuromodulation has also been shown to indirectly include astrocyte signals *in vitro* [18] and auditory signals *in vivo* [19, 20]. To identify the underlying molecular mechanisms, we and others have been examining how ultrasound affects neurons in tractable invertebrate systems [16, 21-23] or mammalian cell [24, 25] and slice cultures [3, 26].

The nematode, *C. elegans* with just 302 neurons connected by identified chemical and electrical synapses generating robust behaviors with powerful genetic tools is ideally suited to probe the molecular effects of ultrasound on neuronal membranes [27-29]. We previously showed that the pore-forming subunit of the mechanosensitive ion channel TRP-4 is required to generate behavioral responses to a single 10 ms pulse of ultrasound generated from a 2.25 MHz focused transducer [21]. This channel is specifically expressed in few dopaminergic (CEPs, and ADE) and interneurons (DVA and PVC) in *C. elegans*, where it has been shown to be involved in regulating the head movement and locomotion [30, 31]. Surprisingly, we found that ectopically expressing this TRP-4 in a neuron rendered that neuron sensitive to ultrasound stimuli, confirming ultrasound-triggered control (Sonogenetics) [21]. Moreover, a second mechanosensitive protein, a DEG/ENaC/ASIC ion channel MEC-4 was also shown to be required for behavioral responses to a 200 ms duration, 1 KHz pulse repetition frequency at 50% duty cycle generated from a 10 MHz focused Ultrasound transducer [22]. These two studies confirm that ultrasound effects on *C. elegans* behavior is likely mediated by mechanosensitive proteins.

In this study, we used genetic tools in *C. elegans* to test whether TRP-4 and MEC-4 act in parallel to mediate the behavioral effects of Ultrasound stimuli. We generated a *trp-4 mec-4* double mutant and compared its ultrasound responses to both single mutants. In addition, we found that ectopically expressing TRP-4 in specific chemosensory neurons can rescue the behavioral deficits in both *trp-4* and *mec-4* null mutants, confirming that these genes act in parallel pathways. Our study demonstrates that multiple mechanosensory pathways act in concert to generate behavioral responses to ultrasound stimuli.

## Results

To test the behavioral responses of *C. elegans* to various ultrasound stimuli, we aligned a transducer with a holder that positioned agar plates at the water level in a tank (**Fig 1A**). Animal responses were captured using a camera and analyzed (**Fig 1B, 1D**, See Methods for more details). Next, we evaluated both pressure and temperature changes at the agar surface for a single 10 ms pulse of ultrasound stimuli of different intensities. We assessed the area on the agar surface which was ensonified by the ultrasound stimuli and found that our system delivered mechanical, but not temperature changes (**Fig 1C**). We found that ultrasound stimuli delivering peak negative pressures greater than 0.75 MPa amplified by 1-10 μm-sized gas filled microbubbles generated robust responses in wild-type (WT) animals. We analyzed these responses and found that animals generated robust increases in their large reversals (events where the head bends twice or more), but not omega bends or small reversals (where the head bends only once) (**Figure 2, Supplementary Fig 1, Supplementary Video S1**). These data are consistent with previous studies showing that *C. elegans* generates dose-dependent responses to ultrasound stimuli [21, 22].

**Figure 1.**
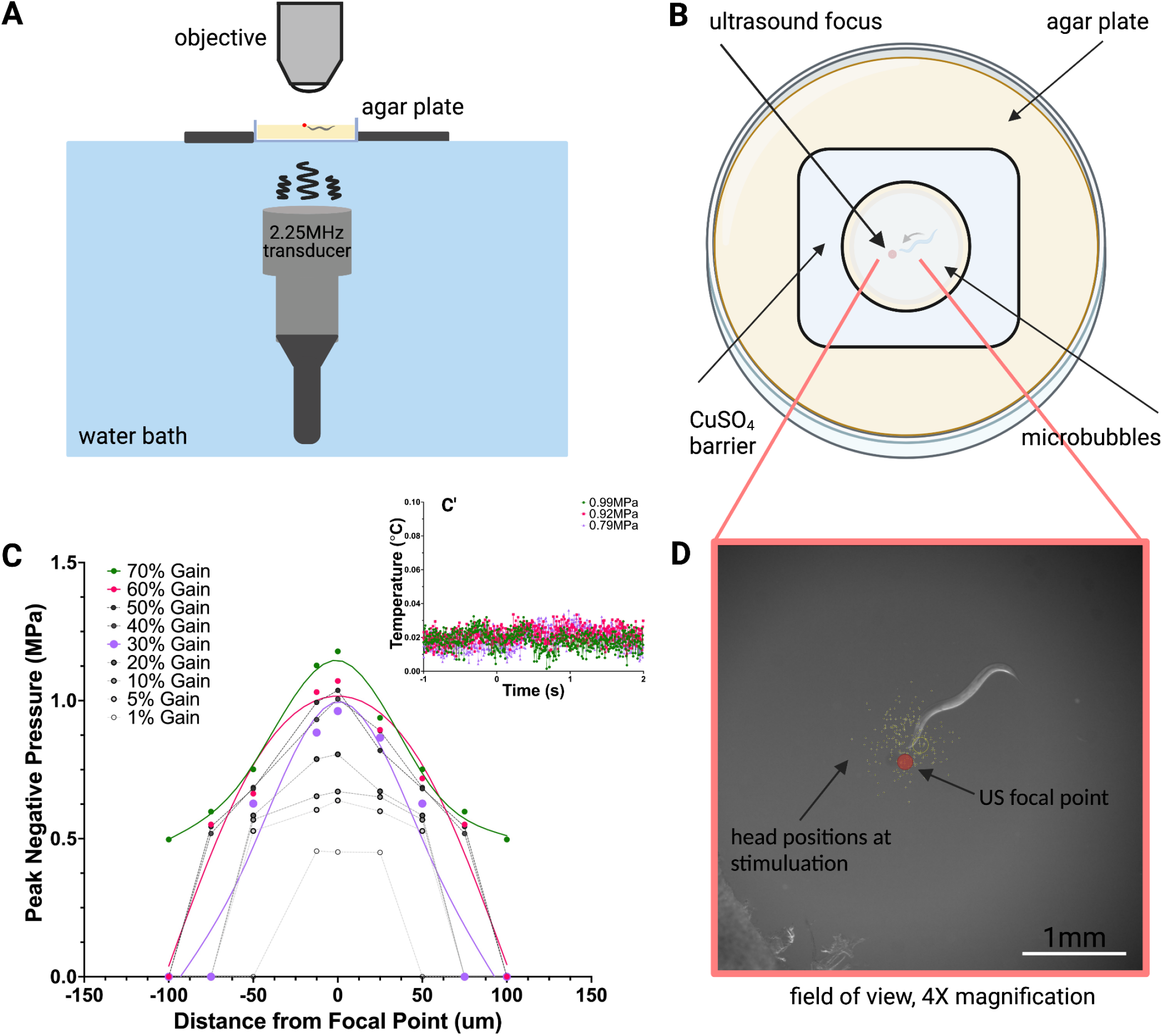
Recording *C. elegans* behavior in response to ultrasound at 2.25MHz. (a) Schematic of 2.25MHz ultrasound imaging system with transducer, water bath, and 4x objective over agar plate. (b) Top view of agar plate with animals corralled by copper sulfate barrier (1.5 cm in diameter) on agar plate with polydisperse microbubbles. (c) Fiberoptic hydrophone measurements at perpendicular distance from focal point of transducer show peak negative pressures ∼1MPa, with (c’) negligible temperature changes at 10ms ultrasound pulses at t=0. Individual points connected via spline fit. (d) Relative head positions (yellow dots) of each animal at time of stimulation.

**Figure 2.**
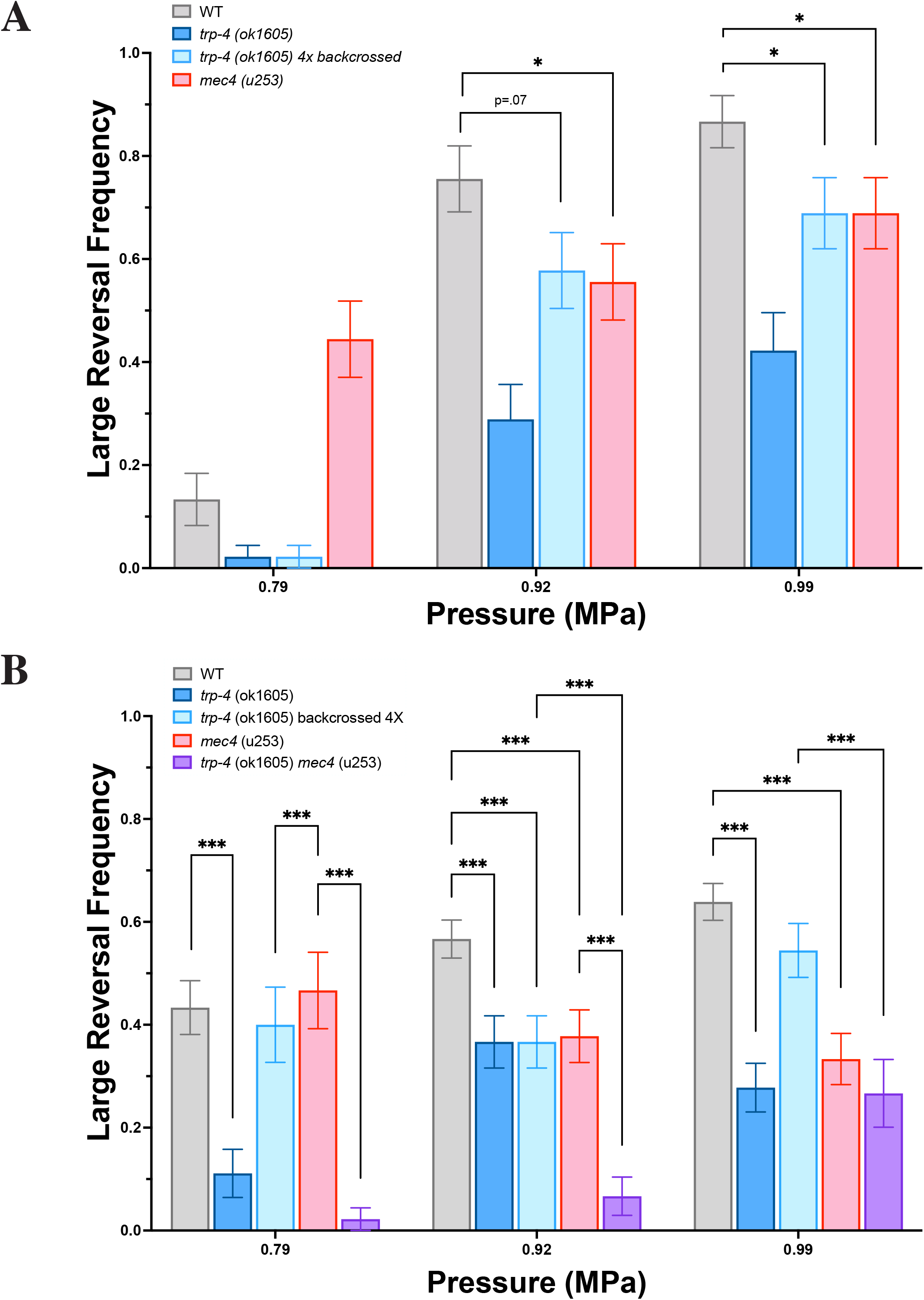
Mutants in mechanosensitive proteins are defective in their responses to ultrasound. Frequency of large reversals (more than two head bends) with and without microbubbles. Large reversal frequency (a) without microbubbles and (b) with microbubbles at various peak negative pressures are quantified. n ≥ 45 for each condition. At certain pressures, the double mutant trp-4 mec4 responds significantly less than each of the individual mutants, which respond less than WT animals. Proportion of animals responding with standard error of the proportion are shown. ***P < .001, **P < .01, *P < .05 by two-proportion z-test with Bonferroni correction for multiple comparisons.

We then analyzed the behavioral responses of null mutants in both the TRP-4 and MEC-4 channels to ultrasound. We found that *trp-4(ok1605)* and *mec-4(u253)* mutants are defective in their ultrasound-evoked large reversal behavior both with and without gas-filled microbubbles (**Fig 2A, 2B**). Moreover, we also found that an outcrossed allele of *trp-4(ok1605 4X)* was also defective in its response to ultrasound stimuli (**Fig 2A, 2B**). This result is consistent with our previous study, which identified a critical role for *trp-4* in mediating ultrasound-evoked behavioral responses [21]. Additionally, we found that *C. elegans* can also respond to ultrasound stimuli even in the absence of gas-filled microbubbles, consistent with a previous study [22]. Moreover, we find that at least at lower ultrasound intensities (0.79 MPa), the probability of *C. elegans’* responses are increased in the presence of microbubbles (**Fig 2A, 2B**). Importantly, we found that the *mec-4(u253) trp-4(ok1605 4X)* double mutant had a stronger defect in ultrasound-trigged large reversal response (**Fig 2B**), suggesting that these genes likely act in parallel. Moreover, we do not observe any consistent change in small reversals and omega bends in any single or double mutants (**Supplementary Fig 1**). Collectively, these data indicate that MEC-4 and TRP-4 mutants might be acting in parallel to mediate *C. elegans* behavioral responses to ultrasound stimuli.

To confirm whether these two mechanosensitive proteins are acting in parallel, we tested combinations of transgenic animals ectopically expressing TRP-4 in various null mutant backgrounds. We expressed TRP-4 under ASH and AWC-chemosensory neuron selective promoters and analyzed the ultrasound responses of the resulting transgenics. While ASH expression of TRP-4 in wild type did not alter its ultrasound-behavior, AWC expression of TRP showed increased frequency of large reversals at 0.79 MPa (**Fig 3A, 3B**). These data are consistent with previous studies, which showed that activation of AWC chemosensory neurons triggers reorientation responses [21, 32]. In contrast, we found that ASH expression of TRP-4 was able to partially rescue behavioral deficits in both the *trp-4(ok1605)* and *mec-4(u253)* null mutants (**Fig 3B**). Moreover, at higher pressures AWC expression of TRP-4 was unable to rescue either single mutant confirming that TRP-4 was likely to selectively function in ASH neurons to drive large reversals in response to ultrasound (**Fig 3B**). Also, we did not observe any consistent changes in small reversals and omega bends (**Supplementary Fig 1**). These data suggest that TRP-4 can function in ASH chemosensory neurons to generate behavioral responses to ultrasound stimuli. Further, these data suggest that TRP-4 and MEC-4 likely act in parallel to generate large reversals in response to ultrasound stimuli.

**Figure 3.**
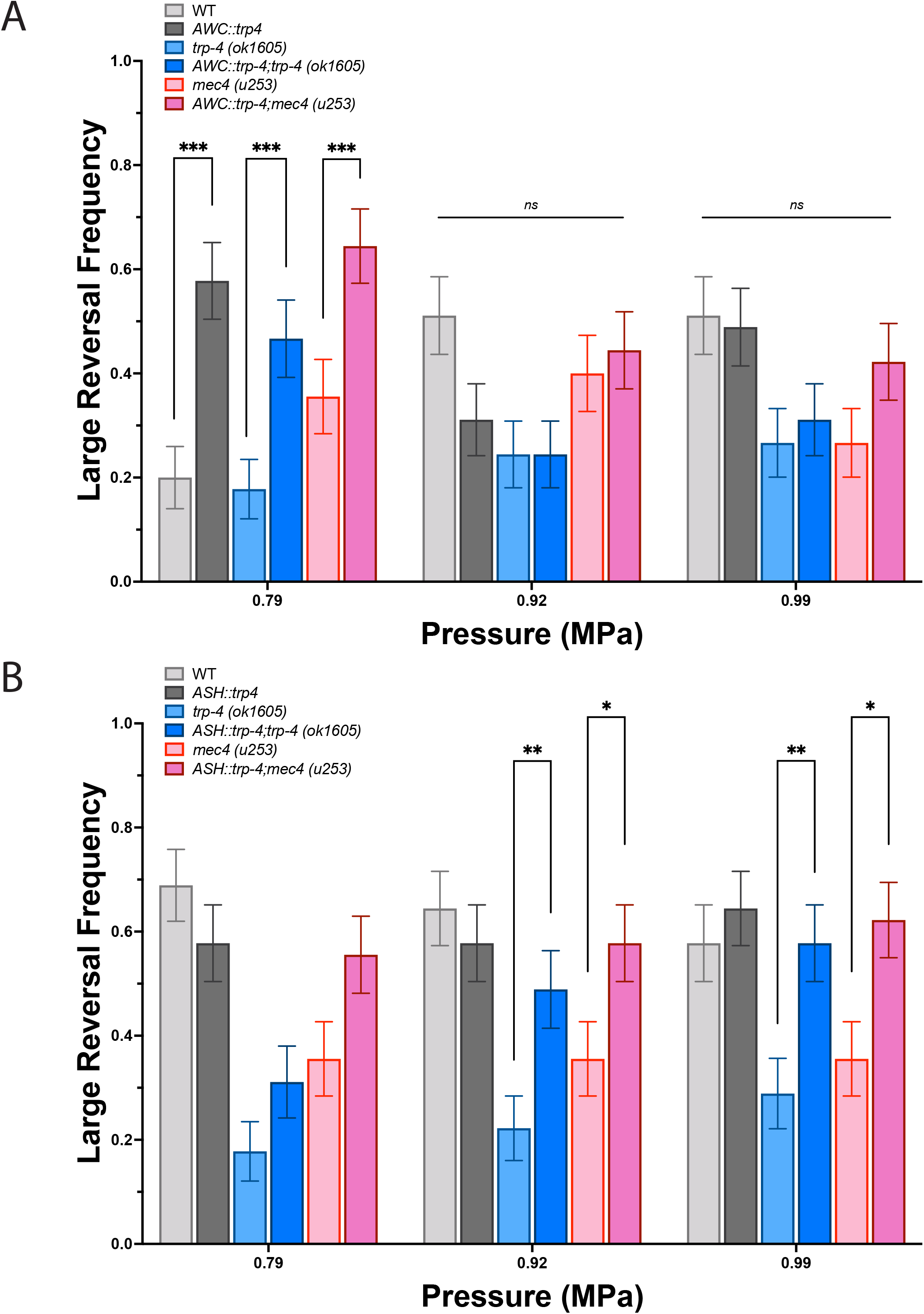
MEC-4 and TRP-4 act in parallel to mediate Ultrasound-evoked behavioral changes. Frequency of large reversals (more than two head bends) in strain ectopically expressing trp-4. Large reversal frequency with (a) AWC::trp-4 and (b) ASH::trp-4 in different backgrounds at different peak negative pressures are quantified. n ≥ 45 for each condition. At certain conditions, trp-4 expression significantly increases large reversal frequency in both trp-4 knockout and mec-4 knockout animals. Proportion of animals responding with standard error of the proportion are shown. ***P < .001, **P < .01, *P < .05 by two-proportion z-test with Bonferroni correction for multiple comparisons.

## Discussion

We showed that a double mutant that deleted both MEC-4 and TRP-4 had stronger behavioral deficits compared to either single mutant alone. Additionally, we showed ASH-specific expression of TRP-4 could rescue the deficit in both single null mutants confirming that these two pathways act in parallel to drive ultrasound-evoked changes in large reversals.

The nematode *C elegans* has provided insights into our understanding of how ultrasound affects animal behavior. We previously showed that ultrasound evoked behavioral changes required the pore-forming subunit of the TRP-4 mechanosensitive channel [21]. This protein is selectively expressed in a few dopaminergic and interneurons and is likely involved in generating head movement and coordinating locomotory behaviors. We suggested that delivering ultrasound to the head of the animal likely activates this channel resulting in reversal behavior [21]. Moreover, a second mechanosensitive channel, MEC-4 (DEG/ENaC/ASIC) has been shown to be required for ultrasound-evoked behavioral responses in *C. elegans* [22]. MEC-4 is a key component of the touch sensitive mechanosensitive ion channel and is expressed in the ALM, PLM, AVM, PVM, FLP and other touch-activated neurons [33, 34]. This study indicated that ultrasound delivered to the head of the animal would also generate a reversal response [22]. While we are unable to replicate this study (**Supplementary Fig 2**), we find that *mec-4(u253)* and *trp-4(ok1605 4X)* mutants are indeed defective under our stimulus conditions and likely act in parallel to generate ultrasound-evoked behavioral changes.

Ultrasound has been used to non-invasively manipulate both neuronal and non-neuronal cells in a number of animals including humans. We show that animals missing two mechanosensitive proteins are defective in their responses to ultrasound stimuli, confirming a role for mechanosensation in mediating the biological effects of this modality. This is also consistent with multiple studies identifying other mechanosensitive proteins that can confer ultrasound sensitivity to mammalian cells *in vitro* and *in vivo* [24-26, 35-37]. This is particularly relevant to the method of using ultrasound to selectively and non-invasively manipulate cells within an animal (“Sonogenetics”). Furthermore, our study implies that ultrasound might affect at least two mechanosensitive ion channels to affect animal behavior. Identifying downstream signaling pathways of these and other ultrasound-sensitive mechanosensitive channels would provide a framework to decode ultrasound neuromodulation and enhance sonogenetic control.

## Methods

### Ultrasound Imaging Assay

A schematic of the system for imaging ultrasound-triggered behavior appears in Figure 1A. An immersible 2.25MHz central frequency point focused transducer (V305-SU-F1.00IN-PTF, Olympus NDT, Waltham, MA) was positioned in a water bath below a 60mm 2% agar-filled petri dish, and connected via waterproof connector cable (BCU-58-6W, Olympus). A single 10ms pulse was generated using a TTL pulse to trigger a multi-channel function generator (MFG-2230M, GWINSTEK, New Taipei City, Taiwan); the amplitude of the signal was adjusted through a 300-W amplifier (VTC2057574, Vox Technologies, Richardson, TX) to achieve desired pressures. *C. elegans* behavior was captured through a high speed sCMOS camera (Prime BSI, Photometrics, Tuscon, AZ) and a 4x objective (MRH00041, Nikon, Chicago, IL). Worm movement was compensated via a custom joystick-movable cantilever stage (LVP, San Diego, CA) controlled by Prior Box (ProScanIII, Prior, Cambridge, UK). All components were integrated via custom Metamorph software (Molecular Devices, San Jose, CA). A goose-neck lamp (LED-8WD, AmScope, Irvine, CA) at ∼45° provided oblique white light illumination.

The ultrasound transducer was focused in the Z-plane to the focal plane of the camera, allowing for X-Y motion of the stage and petri dish using the joystick-movable stage all in the focal plane of both the ultrasound transducer and camera. The petri dish was coupled to the ultrasound transducer via degassed water in the water bath.

For 10MHz experiments, the 2.25MHz transducer was replaced with a 10MHz line-focused transducer (A327S-SU-CF1.00IN-PTF, Olympus) coupled via plastic 20mL syringe and degassed water as previously described^22^. The plastic portion of the petri dish was removed to couple the transducer directly to the agar slab, and a 2x objective (MRD00025, Nikon) replaced the 4x objective to allow for full imaging of the larger focal area of the transducer.

### Behavioral Assays

For experiments with microbubbles, polydisperse microbubbles (PMB, Advanced Microbubbles, Newark, CA) were diluted to a concentration of ∼4×10^7^ and added to an empty 2% agar plate 20 minutes before imaging to allow for absorption/evaporation of the solvent media, leaving microbubbles on the surface of the agar. A dry filter paper with a 1cm hole previously soaked with 200mM copper sulfate solution was placed around the microbubble lawn, and a young adult *C. elegans* was moved from a home plate to the imaging plate using an eyelash. The agar plate was moved around using the motorized stage to place the worm into the focal zone of the transducer where it was stimulated with a single ultrasound pulse of appropriate amplitude. Videos were recorded for 10 seconds at 10 frames/second, with ultrasound stimulation (described above, via TTL pulse) occurring at 1.5 seconds.

Reversals with more than two head bends were characterized as large reversals, those with fewer than two head bends were characterized as small reversals, and omega bends were those which led to a high-angled turn that lead to a substantial change in direction of movement^21^. Wild type animals were tested daily to monitor and maintain a baseline level of reversal behavior, and comparisons between strains were made for animals recorded within same days of testing. Behavioral data were collected over at least three days to confirm reproducibility, the data were then pooled for final statistical analysis, shown in relevant figures.

#### C. elegans

Wild type *C. elegans* – CGC N2; VC1141 *trp4(ok1605);* GN716 *trp4(ok1605)* outcrossed four times (Kubanek et al, 2018); TU253 *mec4(u253)*^22^; IV903 *trp-4(ok1605) mec-4(u253)*, made by crossing GN716 and TU253.

ASH rescues: IV133 *ueEx71 [Psra-6::trp-4; Pelt-2::GFP]* made by injecting N2 with 50ng/μL Psra-6::trp-4, 10ng/μL elt-2::gfp for ASH overexpression of trp-4. IV160 *trp-4(ok1605) I; ueEx88 [Psra-6::trp4; Pelt-2::GFP]* made by injecting VC1141 with 50ng/μL Psra-6::trp4, 10ng/μL elt-2::gfp for ASH rescue of trp-4. IV840 *mec-4(u253) X; ueEx71 [Psra-6::trp-4; Pelt-2::GFP]* made by crossing IV133 and TU253.

AWC rescues: IV157 *ueEx85 [Padr-3::trp-4; Pelt-2::GFP]* made by injecting N2 with 50ng/μL Podr-3::trp4, 10ng/μL elt-2::gfp for AWC overexpression of trp-4. IV162 *trp-4(ok1605) I; ueEx89 [Padr-3::trp-4; Pelt-2::GFP]* made by injecting VC1141 with 50ng/μL Podr-3::trp4, 10ng/μL elt-2::gfp for AWC rescue of trp-4. IV839 *mec-4(u253) X; ueEx85 [Podr-3::trp-4; Pelt-2::GFP]* made by crossing IV157 and TU253.

### Behavioral Ultrasound Pressure and Temperature Measurements

Pressure and temperature measurements were collected through 2% agar plates using a Precision Acoustics Fiber-Optic Hydrophone connected to a 1052B Oscilloscope (Tektronix, Beaverton, OR). The hydrophone probe was moved sub-mm distances using the same stage used for animal recordings while the petri dish was held in place using a three-prong clamp.

### Statistical Analysis

Data were analyzed by combining several days’ recordings within several days’ experimental sessions, with sample sizes chosen to reflect those described previously^22^. All behavioral data were plotted as proportion of response plus/minus standard error of the proportion. For significance tests, two-proportion z-tests were used with Bonferroni corrections for multiple comparisons. All sample sizes were >30, and animals were chosen at random from their broader population. The observer was not blind to the genotype of the group being tested. Animals were excluded from the study if they showed visible signs of injury upon transfer to the assay.

## Acknowledgements

We thank S. Xu, M. Goodman and the CGC for strains. We also thank J. Kubanek for technical advice and A. Singh, and members of the Chalasani and Friend labs for helpful comments and suggestions on the manuscript. This work was funded by grants from the National Institutes of Health (R01MH111534, R01NS115591) (S.H.C.) and from the W.M. Keck Foundation (J.F.),

## Author Contributions

U.M. and C.W. conceived and conducted the experiments, interpreted the data, and co-wrote the paper. A.V., K.C.R., conducted experiments. J.F., interpreted the data and co-wrote the paper. S.H.C. conceived the experiments, interpreted the data and co-wrote the paper. All co-authors provided feedback on the manuscript.

## Supporting Information

**Supplementary Figure 1** | Frequency of small reversals, defined as fewer than two head bends, and omega bends in different strains at different peak negative pressures. (a,b) Small reversals (left) and omega bends (right) from Figure 2 recordings. (c,d) Small reversals (left) and omega bends (right) from Figure 3 recordings. n ≥ 45 for each condition. Proportion of animals responding with standard error of the proportion are shown..

**Supplementary Figure 2** | **Schematic of 10MHz behavioral imaging setup as described in Kubanek et al 2018**. (a) Experimental setup has agar slab with *C. elegans* corralled by a copper sulfate barrier resting on top of a 20mL syringe. Degassed water couples the piezoelectric line-focused transducer (10MHz) to the agar slab. (b) Hydrophone measurements at different perpendicular positions relative to the transducer line focus, with peak negative pressures reaching 1MPa at highest amplifier settings. Yellow bar represents line focus of highest pressure, points connected via spline fit. (c) In both presence and absence of microbubbles, large reversals in WT *C. elegans* are minimal. (d) Example image of *C. elegans* on agar slab approaching ultrasound focal line. Yellow dots represent head positions of each of 224 worms, indicating a significant proportion of ultrasound stimulations occurred when the head was positioned within the high-pressure band (yellow).

**Supplementary Video S1** | **WT *C. elegans* performing a large reversal (and omega bend) in response to ultrasound**. 10 second video recorded at 10fps, with ultrasound pulse at 1.5s (15 frames). Ultrasound label temporally extended for visibility. After ultrasound pulse, worm reverses with >2 head bends and completes an omega bend reorientation.

