## Supplementary figures and images for "Two parallel pathways are required for ultrasound-evoked behavioral changes in *Caenorhabditis elegans*"

### Supplementary Figure S1

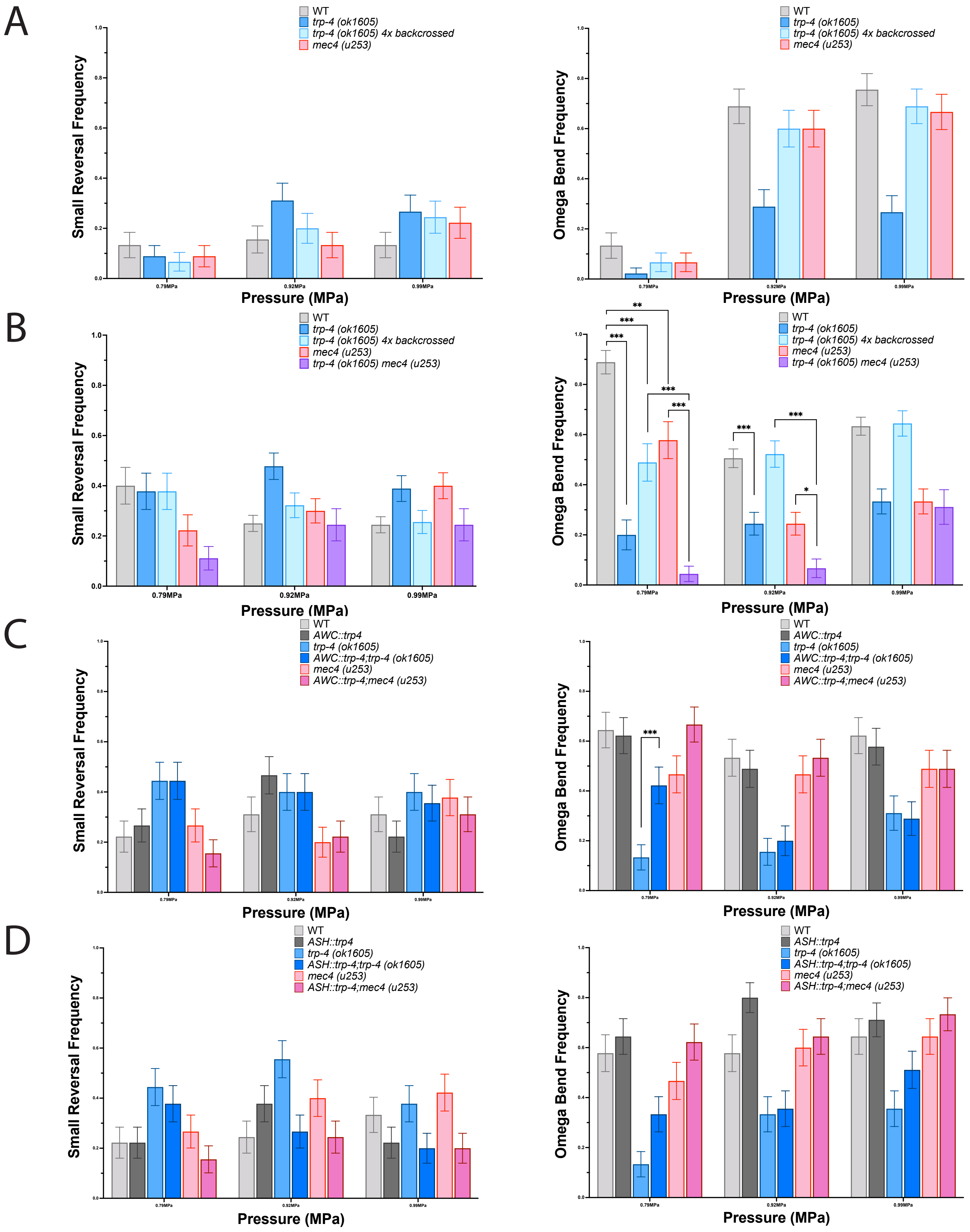

### Supplementary Figure S2

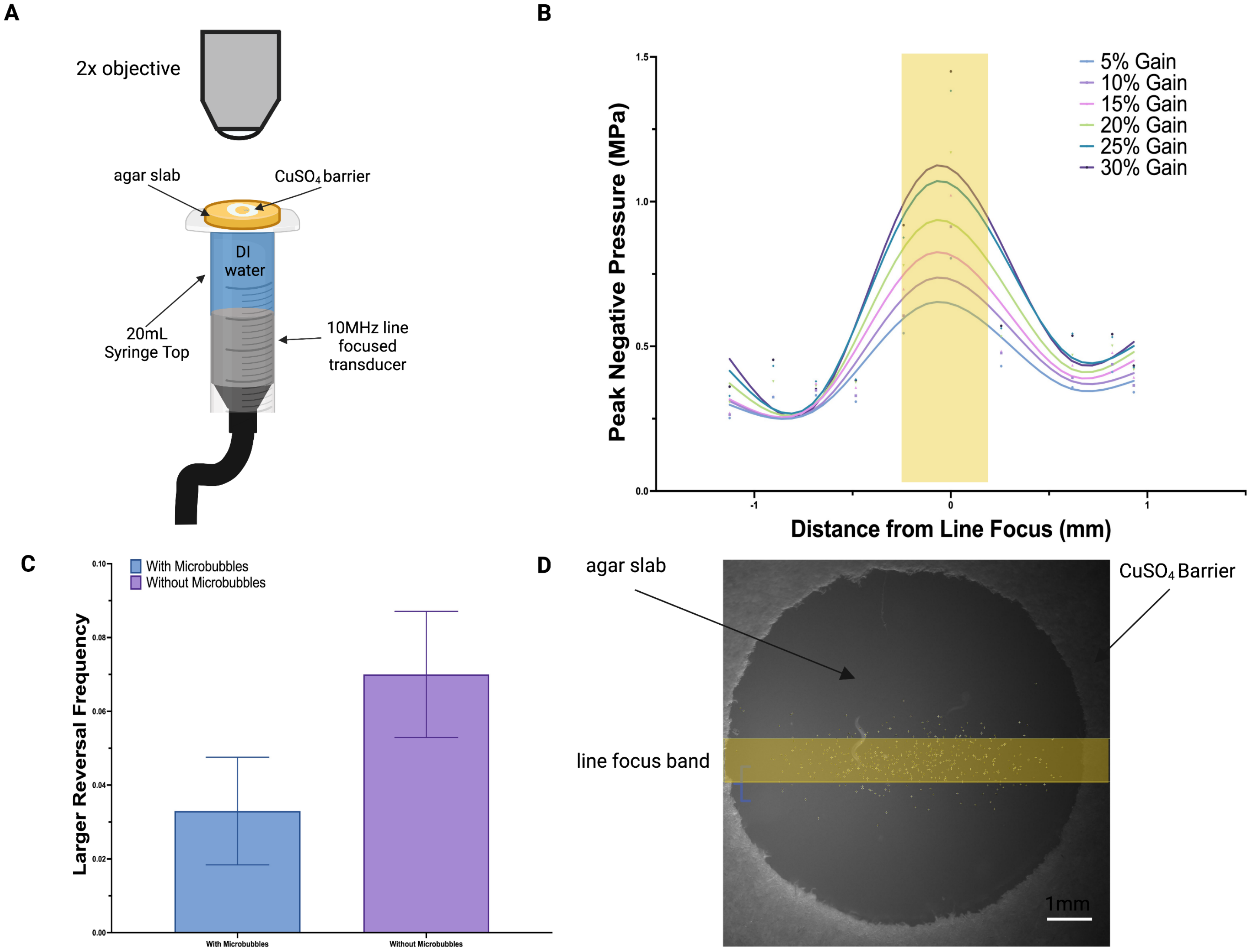
